# A Minimal Biophysical Model of Neocortical Pyramidal Cells: Implications for Frontal Cortex Microcircuitry and Field Potential Generation

**DOI:** 10.1101/2020.01.29.925180

**Authors:** Beatriz Herrera, Amirsaman Sajad, Geoffrey F. Woodman, Jeffrey D. Schall, Jorge J. Riera

**Author notes:** Corresponding author, (JR).

## Abstract

Ca^2+^ spikes initiated in the apical dendrites of layer-5 pyramidal cells (PC) underlie nonlinear dynamic changes in the gain of cellular response, which is critical for top-down cognitive control. Detailed models with several compartments and dozens of ionic channels have been proposed to account for this Ca^2+^ spike-dependent gain with its associated critical frequency. However, current models do not account for all known Ca^2+^-dependent features. Previous attempts to include more features have required increasing complexity, limiting their interpretability and utility for studying large population dynamics. We present a minimal 2-compartment biophysical model, overcoming these limitations. In our model, a basal-dendritic/somatic compartment included typical Na^+^ and K^+^ conductances, while an apical-dendritic/trunk compartment included persistent Na^+^, hyperpolarization-activated cation (I_*h*_), slow inactivation K^+^, muscarinic K^+^, and Ca^2+^ L-type. The model replicated the Ca^2+^ spike morphology and its critical frequency plus three other defining features of layer-5 PC synaptic integration: linear frequency-current relationships, backpropagation-activated Ca^2+^ spike firing, and a shift in the critical frequency by blocking I_*h*_. Simulating 1,000 synchronized layer-5 PCs, we reproduced the current source density patterns evoked by Ca^2+^-spikes both with and without I_*h*_ current. Thus, a 2-compartment model with five non-classic ionic currents in the apical-dendrites reproduces all features of these neurons. We discuss the utility of this minimal model to study the microcircuitry of agranular areas of the frontal lobe involved in cognitive control and responsible for event-related potentials such as the error-related negativity.

**Significance Statement:** A tractable model of layer-5 pyramidal cells replicates all known features crucial for distal synaptic integration in these neurons. This minimal model enables new multi-scale investigations of microcircuit functions with associated current flows measured by intracranial local field potentials. It thus establishes a foundation for the future computational evaluation of scalp electroencephalogram signatures imprinted by Ca^2+^ spikes in pyramidal cells, a phenomenon underlying many brain cognitive processes.

## Introduction

Models of the neocortical microcircuit with biophysically plausible pyramidal cells (PC) are necessary to translate between observed patterns of neural spiking, local field potentials (LFP), and the derived scalp electroencephalogram (EEG). Layer 5 (L5) PCs have an elongated morphology with dendrites spanning all cortical layers; hence their synaptic activity causes laminar current sources (Einevoll et al., 2013; Reimann et al., 2013). The local synchronization of a large population of L5-PCs produces electric potentials that can be measured on the scalp (Hämäläinen et al., 1993; Riera et al., 2012). Integrative features of L5-PCs have suggested their participation in signaling coincident inputs to basal-dendritic/somatic and apical-dendritic regions (Larkum, 2013). The arrival times of sensory inputs, efferent copies, and task rules in agranular frontal cortex are critical in cognitive control (Sajad et al., 2019; Subramanian et al., 2019). One well-characterized cognitive control function is error monitoring by the medial frontal cortex (Stuphorn et al., 2000; Sajad et al., 2019), which is indexed by an error-related negativity (ERN) in scalp potentials (Gehring et al., 1993). Therefore, models of L5-PCs will help clarify the electrogenesis of the ERN.

L5-PCs exhibit two distal excitability zones, endowing these neurons with important integrative features. One excitability zone, at the axon hillock, produces typical Na^+^ action potentials (AP) and another, in the distal trunk, produces Ca^2+^-spikes (Amitai et al., 1993; Yuste et al., 1994; Schiller et al., 1997; Larkum and Zhu, 2002). The coincidence of a Na^+^-AP with an apical dendritic excitatory postsynaptic potential produces additional APs via a backpropagation-activated Ca^2+^-spike, “BAC” firing (Larkum et al., 1999b). Na^+^-APs show a linear frequency-current (f-I) relation with different sensitivities at the two excitability zones (Larkum et al., 2004). Dendritic Ca^2+^-spikes generated by strong inputs show a sustained depolarization (Larkum et al., 2001) that produces high-frequency Na^+^-APs (Schwindt and Crill, 1999; Williams and Stuart, 1999; Larkum et al., 2001). L5-PCs exhibit a critical frequency (CF) between 60 and 200 Hz for eliciting Ca^2+^-spikes (Larkum et al., 1999a) via somatic stimulation, which is sensitive to the hyperpolarization-activated cation current, I_*h*_, in apical-dendrites (Berger et al., 2001).

Previously proposed biophysical models with 2 or 3 neuronal compartments and fewer conductances explained only isolated features (i.e., the I-f curves - Larkum et al., 2004; the BAC-firing - Chua et al., 2015; and Yi et al., 2017). Biophysical models of higher complexity accounted for some combinations of the three major features: the BAC-firing (Rapp et al., 1996; Schaefer et al., 2003; Hay et al., 2011; Bahl et al., 2012; Almog and Korngreen, 2014; Mäki-Marttunen et al., 2018), the f-I curves (Hay et al., 2011; Bahl et al., 2012; Mäki-Marttunen et al., 2018), and the CF of Ca^2+^-spikes (Schaefer et al., 2003; Hay et al., 2011; Bahl et al., 2012; Almog and Korngreen, 2014). However, single cell models with many compartments and ionic channels are computationally expensive to use in large-scale simulations of neocortical networks. Furthermore, fitting these complex models to LFP/EEG data is practically unattainable, limiting interpretability and their applications to other research areas. Only one previous model replicated realistic [Ca^2+^] dynamics in the distal-trunk during Ca^2+^-spikes (Mäki-Marttunen et al., 2018). Furthermore, no previous model has accounted for the I_*h*_ shift of CF, the current source density (CSD) patterns associated with dendritic Ca^2+^-spikes evoked by somatic stimulation of PCs above the CF, and the effect of blocking I_*h*_ on these patterns (Suzuki and Larkum, 2017).

We describe the simplest possible biophysical model (2-compartments, 7 ionic conductances) of L5-PCs accounting for all these features. In particular, it reproduced Ca^2+^ dynamics above the CF and explained the shift produced by I_*h*_. The model replicates CSD patterns obtained from synchronized Ca^2+^-spikes of 1,000 L5-PCs evoked by supra-CF somatic stimulation. Therefore, this minimal L5-PC model will be crucial for the interpretation of LFP-CSD/EEG patterns associated with cognitive control based on our understanding of the agranular laminar microcircuitry (Sajad et al., 2019).

## Materials and Methods

### L5-PC minimal model

We modeled the L5-PC as a 2-compartment neuron with a compartment representing the basal-dendrites/soma, and another compartment representing its distal-trunk (Ca^2+^-spike initiation zone) and the apical-dendrites. The trunk is represented by a transfer resistance (*R*_*T*_) between the two compartments (Figure 1A). The basal-dendritic/somatic compartment includes the classic Hodgkin-Huxley sodium (*I*_*Na*_) and potassium delayed rectifier (*I*_*Kdr*_) currents (Hodgkin and Huxley, 1952). The apical-dendrite/trunk compartment includes persistent *Na*^+^ current (*I*_*Nap*_) (Magistretti and Alonso, 1999), *Ca*^2+^ L-type current (*I*_*CaL*_) (Lytton and Sejnowski, 1991), hyperpolarization-activated non-specific cation current (*I*_*h*_) (Kole et al., 2006), muscarinic K^+^ current (*I*_*M*_) (Adams et al., 1982), and the slow-inactivating potassium current (*I*_*Ks*_) (Korngreen and Sakmann, 2000). The membrane potentials of the two compartments are given by the following coupled differential equations:

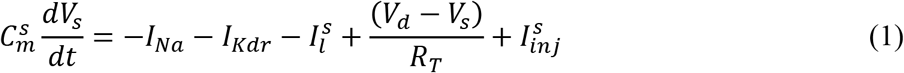

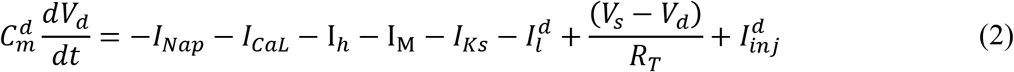

where subscripts *s* and *d* denote the basal-dendritic/somatic and apical-dendritic/trunk compartments, respectively. *V*_*i*_, 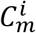, 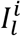 and 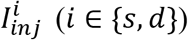 represent the membrane potential, membrane capacitance, leak current and injected current for the *i-th* compartment, respectively (Table 1 – *parameters*). The ionic currents are modeled using the Hodgkin-Huxley formalism in which:

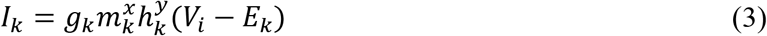

where, *g*_*k*_ is the maximal conductance of the *k-th* ionic channel; *m*_*k*_ and *h*_*k*_ are its activation and inactivation gating variables (Table 2 – *ionic current kinetics*); *x* and *y* are their respective exponents; and *E*_*k*_ is the equilibrium potential of the *k-th* ion. The leak current was modeled by 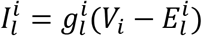. All the equilibrium potentials are considered constant, except for the equilibrium potential of *Ca*^2+^, which depends on the intracellular *Ca*^2+^ concentration ([Ca^2+^]_*i*_) through the Nernst equation. Because of ionic diffusion, we treat [Ca^2+^] as a stochastic variable. Therefore, we added a Wiener noise *g*_*Ca*_*dW*_*Ca*_ to equation (4) using the approach described in a previous study (Riera et al., 2011), with *g*_*Ca*_ = 1 × 10^−9^.

**Figure 1.**
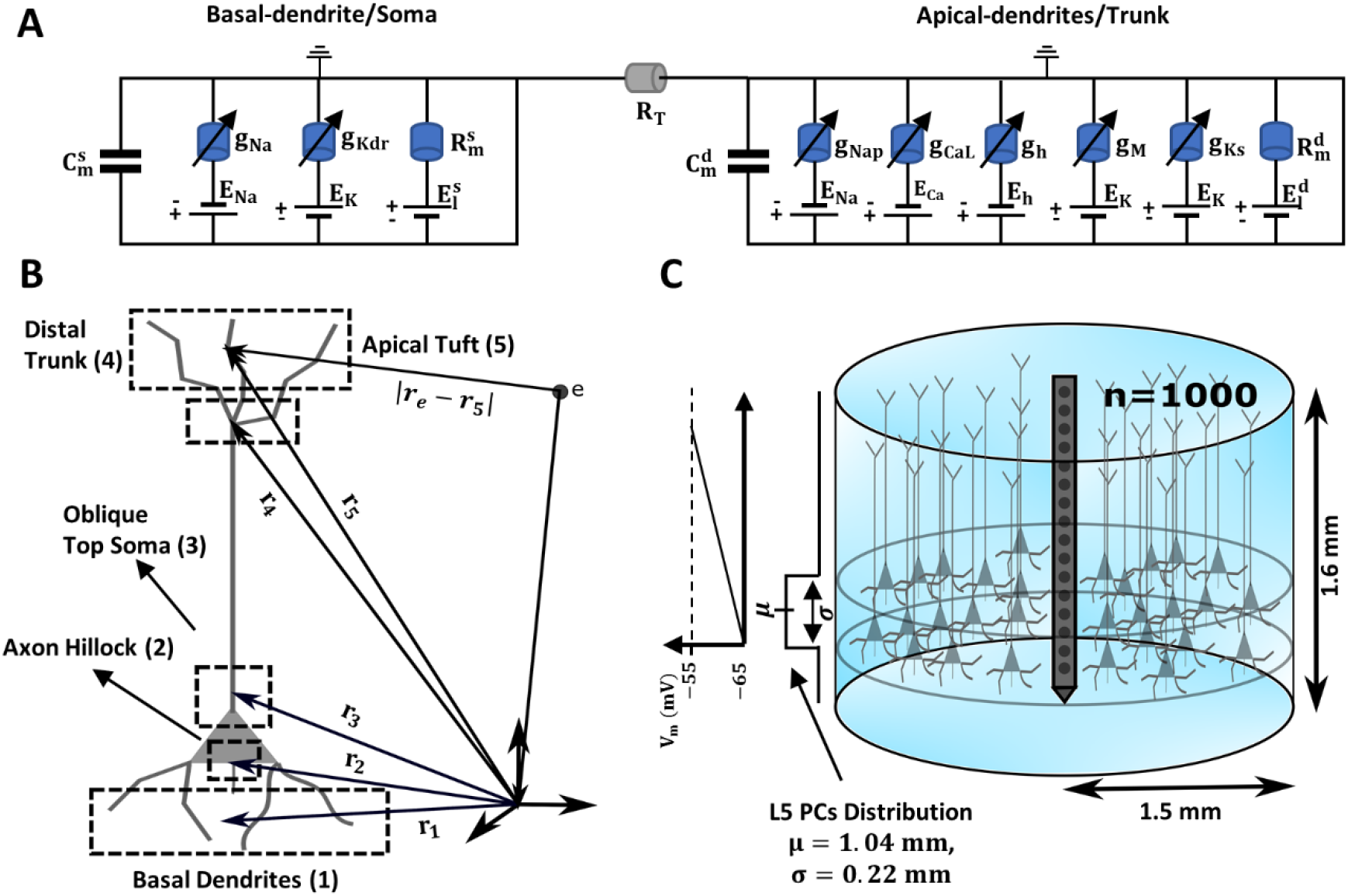
Illustration of the biophysical model, LFP estimation and simulated neocortical column. **A** – Equivalent circuit of the 2-compartment biophysical model. The first and second portions of the circuit represent the basal-dendritic/somatic and apical-dendritic/trunk compartments, respectively. The lengthy trunk is represented by the transfer resistance (*R*_*T*_) between the compartments. Each ionic channel (*k-th*) is represented by an electromotive force ***E***_***k***_ (i.e., the ion equilibrium potential) and a voltage-dependent conductance ***g***_***k***_ in parallel. **B** – Illustration of the forward-modeling used for LFP estimation from the compartmental model of the L5-PCs. To compute the transmembrane currents, the cell was divided into five current source/sink regions (indicated by rectangles). The position of the point source/sink representing the compartment of a neuron is given by the parameter ***r***_***n***_ = {*x*_*n*_, *y*_*n*_, *z*_*n*_}. The position of a representative electrode is given by the parameter ***r***_***e***_ = {*x*_*e*_, *y*_*e*_, *z*_*e*_}. **C** – Diagram of the simulated cortical column formed by a collection of 1,000 L5 PCs. The somas were distributed randomly in the tangential dimension of layer 5. The mean (standard deviation) depth of the neurons was 1.04 (0.22) *mm* from the pia matter. The simulated cortical column had a diameter of 3 *mm* and a total depth of 1.6 *mm*. As previously reported (Berger et al., 2001), we used a resting membrane potential of −65 *mV* at the somatic compartment, which drifted to −55 *mV* at the apical-dendritic/trunk compartment due to the presence of the I_*h*_ current.

**Table 1:** Parameters used for the simulations. The first column indicates the ionic channels per compartment. The second and third columns show the maximum conductance and equilibrium potential for each ionic channel, respectively. The exponents of the activation (x) and inactivation gating are indicated in the fourth column. Electrotonic parameters (capacitances/resistances) are also shown.

| | $g_k$ ($\mu S$) | $E_k$ (mV) | (x,y) |
| --- | --- | --- | --- |
| **Somatic Compartment** |  |  |  |
| $C_m^s = 0.26 \text{ nF}$, $R_m^s = 50 \text{ M}\Omega$ (Larkum et al., 2004) | | | |
| **Na** | 18 | 50 | (3,1) |
| **Kdr** | 5 | -85 | (0,4) |
| **Leak** | $1/R_m^s$ | -31.5 | (0,0) |
| **Dendritic Compartment** |  |  |  |
| $C_m^d = 0.12 \text{ nF}$, $R_m^d = 43 \text{ M}\Omega$ (Larkum et al., 2004) | | | |
| **Nap** | 0.022 | 50 | (3,1) |
| **CaL** | 3.85 | $\frac{RT}{zF} \ln \left( \frac{[Ca^{2+}]_o}{[Ca^{2+}]_i} \right)$ | (2,0) |
| **h** | 0.865 | -45 | (1,0) |
| **M** | 1 | -85 | (1,0) |
| **Ks** | 28 | -85 | (2,1) |
| **Leak** | $1/R_m^d$ | -48.1 | (0,0) |
| $R_T = 65 \text{ M}\Omega$ (Larkum et al., 2004) | | | |
| $[Ca^{2+}]_o = 2 \text{ mM}$ | | | |

**Table 2:**
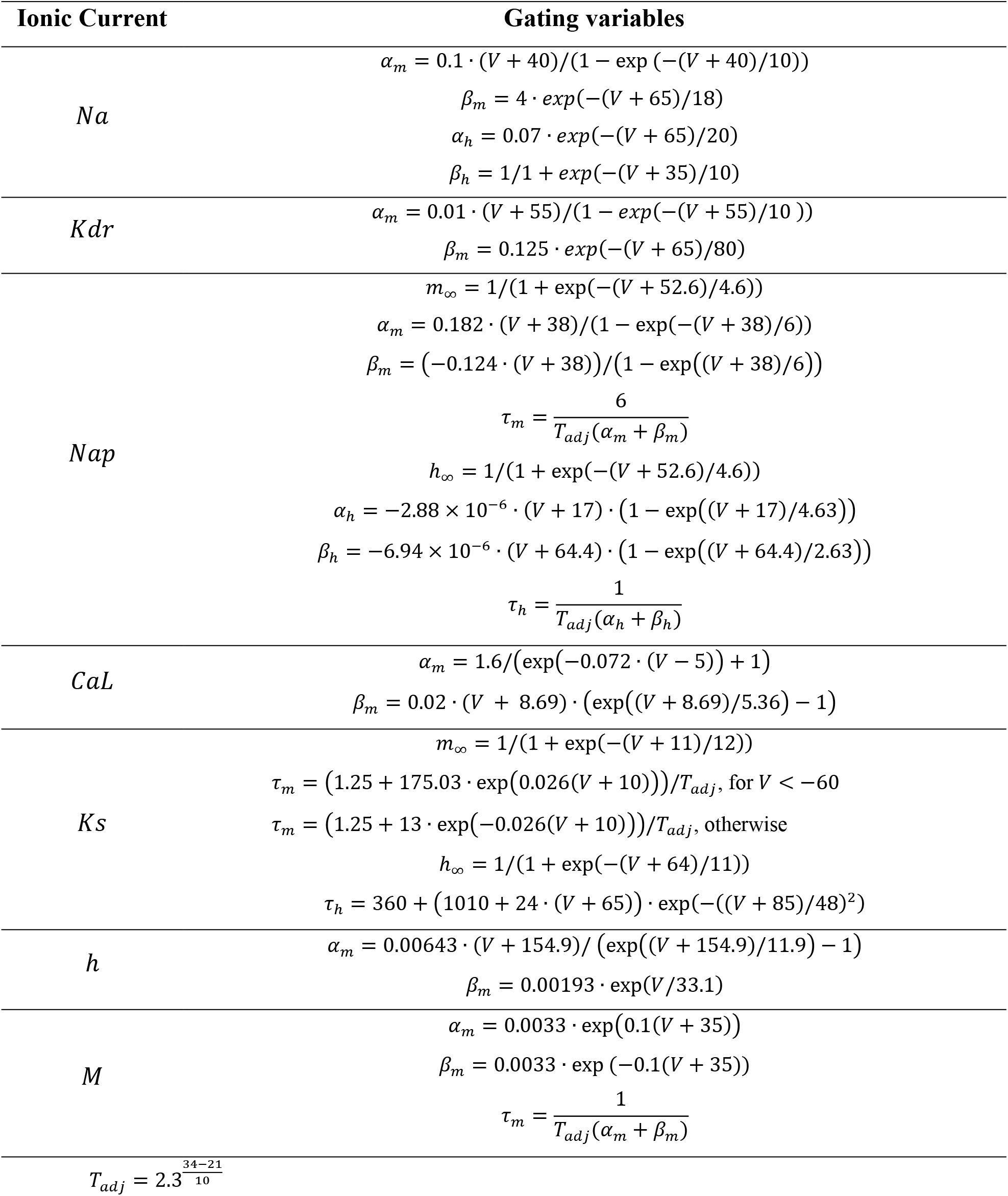
The gating kinetics for each ionic channel.

The intracellular *Ca*^2+^ concentration dynamics is given by

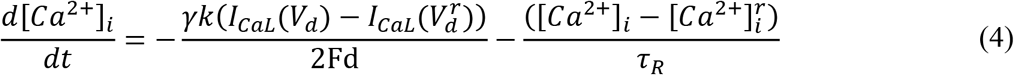

where 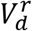 is the dendritic resting potential, 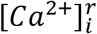 is the intracellular *Ca*^2+^ concentration at rest, *τ*_*R*_ = 80*ms* is the decay time constant of the intracellular *Ca*^2+^ concentration due to active transport (Schaefer et al., 2003). *d* = 1 *μm* is the depth of the submembrane Ca^2+^ shell, *F* = 96489 *C*/*mol* is the Faraday’s constant, and *k* = 10,000/*A*_*d*_ is the unit conversion constant for *I*_*CaL*_ (mA). The surface area *A*_*d*_ = 9302.3 *μm*^2^ of the apical-dendrite/trunk compartment was calculated based on the values given by Larkum et al. (2004) for parameters 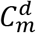 and 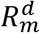. *γ* represents the fraction of free Ca^2+^ (not buffered), which was adjusted to reproduce experimental data for [*Ca*^2+^]_*i*_ in the distal-trunk (Larkum et al., 1999a). The basal intracellular Ca^2+^ was set at its typical physiological value 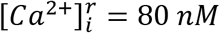.

### Frequency-current (f-I) relation

We create the frequency-current (f-I) curves by injecting a noisy staircase current into either compartment and calculating the somatic firing rate for each current step. The noisy input current was an Ornstein-Uhlenbeck process (Larkum et al., 2004):

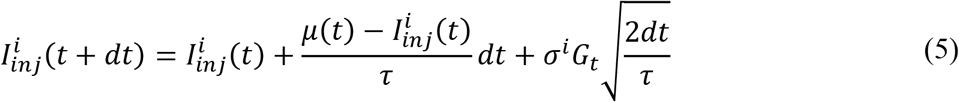

where 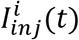 is the injected current at the *i-th* compartment with mean *μ*(*t*), compartment-dependent standard deviation *σ*^*i*^, and time correlation length *τ*. *G*_*t*_ is a random number generated at each time point from a Gaussian distribution with mean = 0 and standard deviation = 1. We set *τ* = 3 *ms* as in the experimental study (Larkum et al., 2004), and *dt*, the time increment, equal to the integration time step. The mean *μ*(*t*) increased over time between 0.2 and 0.75nA as a staircase function with steps of *μ*(*t*) = 0.05*nA* every 2 s.

### Modeling a population of L5-PCs

In addition to reproducing all main features of PC reported from intracellular recording studies, we validated its usefulness to model large-scale extracellular electric potentials (e.g., LFP) generated by cortical microcircuits. To that end, we simulated a neocortical column comprised of 1,000 L5-PCs. For now, they were not connected to each other. Nevertheless, this approach allowed us to determine the transmembrane ionic current densities (active/returning) and laminar LFP associated with synchronized apical-dendritic L5-PC Ca^2+^-spikes. The laminar LFPs and CSD patterns were compared with those obtained by Suzuki and Larkum (2017).

### Calculating the LFPs

We calculate the LFP from the transmembrane currents generated by a collection of neurons using the point-source approximation (Holt and Koch, 1999), which assumes that the transmembrane currents through a compartment can be approximated as a single monopolar source/sink placed in an extracellular medium at the center of the compartment. To compute the transmembrane currents, we divided each compartment into regions (Figure 1B). This approach permits the spatial separation of active ionic and passive returning (i.e., capacitive and leak) currents. The basal-dendritic/somatic compartment was modeled by three regions: the basal dendrites, the axon hillock, and the soma-oblique dendrites. The apical-dendrite/trunk compartment was modeled by two regions: the distal trunk (including the main bifurcation point), and the tufted apical-dendrites. Each region was represented by a single monopolar current source/sink. The ionic and capacitive/leak currents are distributed between these regions as follow:

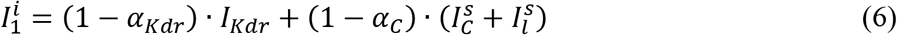

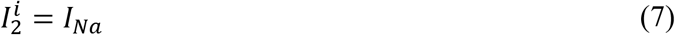

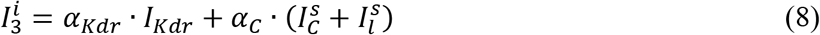

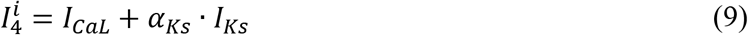

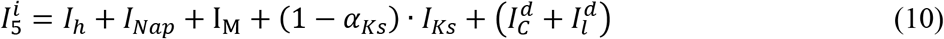

where 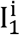, 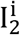, 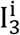, 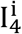, and 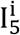 are the total transmembrane currents (Figure 1B) of the basal 1 2 3 4 5 dendrites (1), axon hillock (2), top soma-oblique dendrites (3), distal trunk (4), and the apical-dendrite (5) regions, respectively. 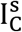 and 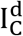 are the somatic and dendritic capacitive currents, respectively; and are equal to 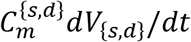. The distribution of ionic currents in these five regions was determined by taking into consideration the following physiological/morphological characteristics.

First, because of their extensive surface area, returning (i.e., capacitive and leak) currents were distributed only in dendrites. The scaling factor *α*_*Kdr*_, *α*_*C*_, and *α*_*KS*_ were adjusted to reproduce the CSD patterns reported by Suzuki and Larkum (2017). We separated the somatic capacitive and the *I*_*Kdr*_ currents into their contribution by the basal and top soma-oblique dendrites. These regions possess a bigger area and a combined higher density of *I*_*Kdr*_ channels than the axon hillock (Ramaswamy and Markram, 2015). In the axon hillock, we included only the Na^+^ current because its density in this area is at least 50-fold higher than at proximal dendrites (Ramaswamy and Markram, 2015). The *I*_*CaL*_ and *I*_*Ks*_ currents were incorporated in the main bifurcation point of the trunk since this region is the Ca^2+^-spike excitability zone (Larkum et al., 1999b). The I_*h*_ current was added to the apical dendrite compartment because of its high density in this region (Kole et al., 2006) and critical influence on synaptically evoked activity in the distal apical dendritic arbor (Harnett et al., 2015). The *I*_*Nap*_ (Schwindt and Crill, 1995) and I_*M*_ (Hay et al., 2011) currents were also included in this area because of their role in the amplification/attenuation of synaptic currents in the distal apical-dendrites. Finally, the capacitive current was added into this region since the distal dendritic arbor covers a greater area than the Ca^2+^-spike excitability zone (Ramaswamy and Markram, 2015).

We compute the LFPs at 16 equally spaced vertically aligned points to simulate the linear microelectrode array (Michigan probe) used by Suzuki and Larkum (2017). As in their study, the inter-electrode distance (ℎ) was 100 *μm*. Motivated by their stimulation protocol with the right-angled prism, we consider that the linear probe was located at the center of a cylindrical neocortical column of 3 *mm* in diameter, and with constant and isotropic electrical conductivity *σ* = 0.323 *S*/*m* (i.e., average across layers from Goto et al., (2010)) (Figure 1C). Given the maximal current produced by individual PCs, 1,000 L5-PCs were required to generate CSD amplitudes in the range reported by Suzuki and Larkum (2017). The electric potential at electrode position 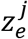 is given by (Nicholson and Llinas, 1971):

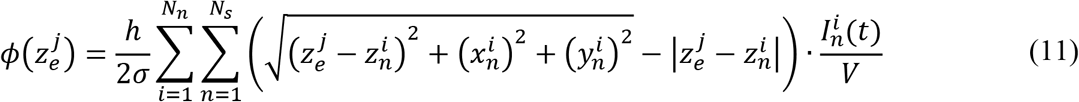

where 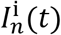 is the transmembrane current generated by the point-source *n* of the neuron *i*; 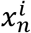, 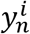 and 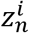 are the coordinates of the point-source *n* of the network neuron *i*, and *V* is the volume of the cortical column. *N*_*s*_ = 1,000 and *N*_*s*_ = 5 represent the total number of neurons in the network and the total number of regions in each neuron, respectively. The 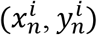 coordinates of the neurons in the simulated neocortical column were generated randomly from a uniform distribution. The 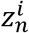 coordinate of the axon hillock point-source/sink of the network neurons was also generated randomly from a uniform distribution with values between 1.025 mm and 1.450 mm (below the pia matter, Suzuki and Larkum (2017)). The location of the basal-dendrite, trunk main bifurcation point and apical-dendrite point-sources were calculated relative to the location of the neurons’ axon hillock. The basal dendrites point-source was always 0.15 mm bellow the axon hillock, the main bifurcation point of the trunk was always 0.89 mm above the axon hillock (Ledergerber and Larkum, 2010, Figure 12), and the apical dendrite point-source was 0.15 mm above the trunk main bifurcation point. The position of the top-soma oblique dendrites, representing part of the somatic returning currents, was generated randomly with values between 0.7 and 1 mm from the cortical surface. The proposed distribution of point-sources for dendrites was inspired by morphological data of L5-PCs (Mohan et al., 2015). Wiener noise *g*_*k*_*dW*_*k*_ was added to equations (1) and (2) to instantiate variability in the timing of L5-PC Na^+^-APs and Ca^2+^-spikes, with *g*_*s*_ = 0.05 and *g*_*d*_ = 0.025.

### Current source density (CSD) analysis

We estimated the CSD patterns evoked by the simulated LFPs using the spline inverse CSD method (spline iCSD) (Pettersen et al., 2006). The iCSD methods are based on the inversion of the solutions of the electrostatics forward problem and assume cylindrical confined and symmetric CSDs. Specifically, the spline iCSD method assumes a continuously varying CSD along the recording electrodes, which is calculated by interpolating a set of cubic splines, requiring the CSD and its first and second derivatives in the vertical direction to be continuous (Pettersen et al., 2006). It also considers a homogeneous disc distribution in the in-plane (*x*, *y*) directions. In agreement with pthe revious section, a homogeneous and isotropic volume conductor with extracellular conductivity of *σ* = 0.323 *S*/*m* (Goto et al., 2010) was used. Based on L5-PC density and the CSD peak amplitudes in Suzuki and Larkum (2017), the diameter of the cylindrical source model was set to 3 *mm*. The estimated CSD based on the simulated LFPs were convolved with a Gaussian filter of *σ* = 0.1 *mm* to produce a spatially smoothed CSD estimate.

### Simulations and code accessibility

Simulations were performed in MATLAB (R2018b, MathWorks) with custom-written scripts. The model equations are solved using the SDETools toolbox for the numerical solution of stochastic differential equations (https://github.com/horchler/SDETools), with a time-step of 1 *μs*. All simulation parameters are listed in Table 1 with ionic channel kinetics in Table 2. To calculate the CSD, we created customized scripts that use the functions provided in the CSDplotter toolbox (https://github.com/espenhgn/CSDplotter), which implements the iCSD methods described in Pettersen et al., (2006). The MATLAB scripts of the model implementation as well as for the LFPs and CSD calculations are publicly available at (https://github.com/beaherrera/2-compartments_L5-PC_model).

## Results

### Model testing approach

Traditionally, parameter estimation of L5-PC biophysical models is performed using quantitative strategies aimed at numerically minimizing model prediction errors while reproducing transmembrane potential traces in specific experimental paradigms. In some cases, the data are used to fit channel kinetics (Rapp et al., 1996), while in others (Hay et al., 2011; Bahl et al., 2012; Almog and Korngreen, 2014; Chua et al., 2015; Mäki-Marttunen et al., 2018) conductance ranges are fitted with generic optimization methods, based on known channel kinetics. However, such a quantitative approach is very challenging if biophysical models are used to simultaneously fit data from multiple experimental paradigms. In such cases, a qualitative trial/error approach based on electrophysiological knowledge about the effect that each ion channel produces on the data is more effective (Schaefer et al., 2003; Larkum et al., 2004; Yi et al., 2017). We will use the qualitative trial/error approach as our goal is to satisfice qualitatively and not satisfy quantitatively six different properties of L5-PCs (Table 3), which were reported using a variety of experimental paradigms. We also employed previously known channel kinetics. The rationale used to determine ionic distributions and conductances is now explained.

**Table 3:** Summary of previous/current biophysical models used to describe the principal features of L5-PCs. The first, second, third and fourth columns show the study, number of compartments, number of ionic channels and the platform used to create the simulated data for each study. The fifth column list the features that were explained by each study. The column “Ions” provides the following information *N*_*s*_: *N*_*c*_/*C*, where *N*_*s*_ and *N*_*c*_ are the number of ionic species considered and the number of ionic channels per compartment, respectively. In some cases, the number of ionic channels per compartment depends on the regions of the neuron considered. Acronyms: IF – Integrated and Fire Model; S – Soma; AD – Apical Dendrites; BD – Basal Dendrites; T – Trunk; and A_h_ – Axon Hillock.

| Study | Geometry | Compartment | Ions | Platform | Features Explained |
| --- | --- | --- | --- | --- | --- |
| (Rapp et al., 1996) | Realistic | >>1,000 | 2: 2/c<br>Total >>2,000 | NEURON | BAC firing |
| (Larkum et al., 2004) | Simplified | 2 | 2: 1/c, +IF model<br>Total = 2 | Not reported | f-I curves |
| (Schaefer et al., 2003) | Realistic | 8 | 7: 6/c<br>Total = 48 | NEURON | BAC firing<br>CF |
| (Hay et al., 2011) | Realistic | 199 | 8: 8/c S(1),<br>6/c AD(198)<br>Total = 1,196 | NEURON | BAC firing<br>CF<br>f-I curves |
| (Bahl et al., 2012) | Realistic | 14 | 8: 1/c A<sub>h</sub>(5), 5/c S(1),<br>1/c BD(1), 4/c AD(7)<br>Total = 39 | NEURON | BAC firing<br>f-I curves |
| (Almog and Korngreen, 2014) | Realistic | Many<br>Compartments<br>50 $\mu m$ in length | 8: 8/c<br>Total = Many | NEURON | BAC firing<br>CF |
| (Chua et al., 2015) | Simplified | 3 | 1: 1/c AD,<br>+ IF model<br>Total = 1 | NEST | BAC firing |
| (Yi et al., 2017) | Simplified | 2 | 3: 2/c S(1), 1/c AD(1)<br>Total = 3 | MATLAB | f-I curves |
| (Mäki-Marttunen et al., 2018) | Realistic | 4 | 10: 9/c S(1),<br>1/c BD(1), 7/c AD(2)<br>Total = 24 | NEURON | BAC firing<br>f-I curves<br>[Ca<sup>2+</sup>]<sub>i</sub> |
| This study | Minimum | 2 | 7: 2/c BD/S(1),<br>5/c AD/T(1)<br>Total = 7 | MATLAB | BAC firing<br>CF<br>f-I curves<br>[Ca<sup>2+</sup>]<sub>i</sub><br>I<sub>h</sub> effect<br>CSD maps |

Ion channels for each compartment were selected based on experimental findings and modeling studies. In the soma, we included the classic Na^+^ and K^+^ delayed rectifier channels to generate the APs (Hodgkin and Huxley, 1952). Previous studies (Lytton and Sejnowski, 1991; Larkum et al., 2004; Hay et al., 2011; Mäki-Marttunen et al., 2018) reported the need for the after-hyperpolarization (AHP) current to reproduce the f-I relationship shown experimentally by the L5-PCs. However, as in Bahl et al., (2012), this current was not needed to explain the f-I relationship. On the other hand, the *I*_*CaL*_ (Almog and Korngreen, 2009; Pérez-Garci et al., 2013), the *I*_*Nap*_ (Schwindt and Crill, 1995; Crill, 1996) and the *I*_*Ks*_ (Harnett et al., 2013) currents were inserted in the dendritic compartment to generate the characteristic shape of dendritic Ca^2+^-spikes and in agreement with experimental data. *I*_*CaL*_ defined the amplitude and velocity of the initial depolarization phase. *I*_*Nap*_was responsible for the long plateau-like depolarization characteristic of these spikes, and also essential for the CF effect. *I*_*Ks*_ defined both the duration of the repolarization phase and the amplitude of the Ca^2+^-spike’s after-hyperpolarization phase. The I_*h*_ current (Kole et al., 2006) was included in the apical-dendrite/trunk compartment to account for the increase in the resting potential reported in the dendrites of these neurons (Berger et al., 2001). Moreover, this current plays a significant role in the BAC firing modulation and the synaptic integration, as well as in the reported changes in both the CF for Ca^2+^-spike generation (Berger et al., 2003) and the CSD pattern evoked by dendritic Ca^2+^-spikes (Suzuki and Larkum, 2017). Finally, the M-current was needed for the spike repolarization phase when staircase input currents were applied to the apical dendrites. Without this current, the dendritic membrane potential could not complete the repolarization phase. The voltage dependence of the channel kinetics at the apical-dendrite/trunk compartment was shifted by +8 mV to account for the shift in the resting membrane potential.

Henceforth, we tested our L5-PC biophysical model in two steps. We first <u>validate</u> the minimal model by reproducing all known Ca^2+^-dependent synaptic facilitation features. We next assess the capabilities of the model to <u>reproduce</u> the large-scale Ca^2+^-spike dependent LFPs associated with the synchronized activation of a population of L5-PCs in a neocortical column responding to supra-CF somatic stimulation.

### Validation of the model

#### Frequency-current (f-I) relationship

We first investigated whether our model predicts the f-I relationship previously reported for L5-PCs when either the soma or the distal-trunk region is stimulated (**Figure 2**A, Larkum et al., (2004)). We injected into the soma or the distal-trunk, a staircase incrementing noisy input current generated using the Ornstein–Uhlenbeck method (see Materials and Methods), with standard deviation *σ* = 0.2 *nA*, or *σ* = 0.09 *nA*, respectively. Figure 2B shows the somatic AP response (blue, *top panel*) to the somatic input current (blue, *second panel*). Then, the mean somatic AP frequency was computed for each current step. Mean and standard errors of the mean (SEM) over 50 trials were estimated (Figure 2C, blue). Overall, the model predicted a linear f-I relationship for the somatic input current (dashed blue line, goodness-of-fitting R^2^ = 0.959) that fell within the range of two experimentally reported studies for L5 PCs (Figure 2C, black traces/shadow) (Larkum et al., 2004; Bahl et al., 2012). Figure 2B shows the somatic AP response (blue, *third panel*) to the dendritic input current (red, *bottom panel*), and Figure 2C compares observed with simulated values. The model also predicted a linear f-I relationship for dendritic input current (dashed red line, goodness-of-fitting R^2^ = 1.00). In agreement with experimental data (Larkum et al., 2004), current injections at the trunk must be ~300 *pA* larger than those needed at the soma to produce the same AP frequency in these L5-PCs. This effect was quantified using parameter ΔI (Figure 2D), which was calculated for all somatic AP rates from simulated (N = 6, ΔI = 0.3142 ± 0.0140 *nA*) and experimental (N = 6, ΔI = 0.3333 ± 0.0258 *nA*) data. This difference was statistically insignificant (t(5) = 2.0789, p = 0.0922, two-tailed paired t-test) demonstrating the model represents adequately the experimental f-I relationships. However, our model predicted a threshold for somatic AP initiation of ~0.35 *nA* at trunk current injection sites, which was smaller than that of ~0.5 *nA* reported experimentally. This negative finding could be explained by the difference in the injection site along the trunk of the actual L5-PCs used in the experiments (Larkum et al., 2004). Current injections at sites relatively distant to the trunk bifurcation site will require larger amplitudes as the density of CaL channels might be lower. In our model, the mimicked injection was consistently performed at the level of that bifurcation lowering the threshold required to achieve somatic AP firing.

**Figure 2.**
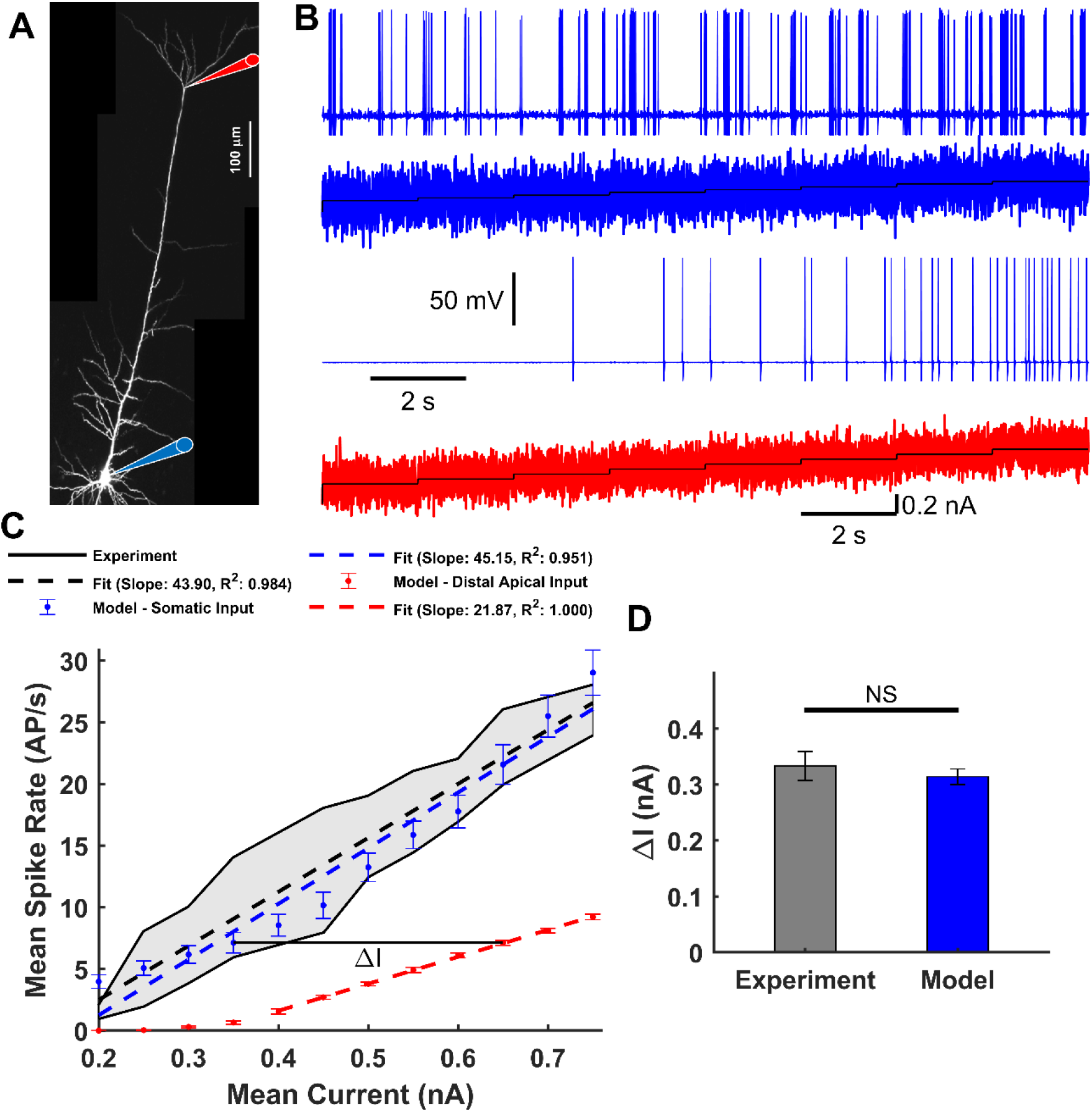
Frequency - Input (f-I) relationship. **A** – Micrograph of a L5-PC with recording locations at the soma (*blue*) and distal trunk (*red*) indicated with diagram pipettes. **B** – Somatic AP responses (1^st^ and 3^rd^ panels) to the staircase incremented noisy input current (2^nd^ and 4^th^ panels) injected into the soma (*blue*) and distal trunk (*red*). **C** – Observed (*black*, Larkum et al., 2004; Bahl et al., 2012) AP firing frequency as a function of the mean input current. The range of observed values is highlighted by a gray fill. Simulated mean and SEM spike rate over 50 trials for each current step in the soma (*blue*) or distal trunk (*red*) compartment. Superimposed are observed (*black dashed*) and simulated (*blue and red dashed*) linear regressions. **D** – Current differences (ΔI) between the f-I curves for somatic and distal trunk stimulation to produce the same Na^+^-AP firing frequency. No significant differences were found between the observed ΔI, numerically estimated from Larkum et al., (2004) and that predicted by the model (t(5) = 2.0789, p = 0.0922, paired two-tailed t-test).

#### Back-propagating AP activated Ca^2+^-spike (BAC) firing

Next, we examined how the L5-PC biophysical model responds and integrates inputs into the distal-trunk and soma (**Figure 3**A) at different times. Firstly, we stimulated the distal-trunk with a subthreshold current generated from a double exponential function of the form (1 − exp(−1/*τ*_2_)) ∙ exp (−1/*τ*_2_) with *τ*_1_ = 2 *ms* and *τ*_2_ = 10 *ms*, and an amplitude of 0.29 *nA*. In agreement with experimental studies (Larkum et al., 1999b; Schaefer et al., 2003), only a small somatic and apical-dendritic/trunk depolarization were evoked by this current injection (Figure 3B). Second, we injected a threshold current pulse (duration: 5 *ms*, amplitude: 1 *nA*) into the soma, which elicited an AP that propagated back to the apical-dendrite/trunk compartment creating a dendritic depolarization but no Ca^2+^-spike (**Figure 3**). Third, we tested the model response when both stimuli were combined. We applied the somatic current pulse and 1 *ms* later the subthreshold current at the trunk. This resulted in the generation of an AP, a dendritic Ca^2+^-spike, and another somatic AP following the onset of the dendritic Ca^2+^-spike (Figure 3D). We could also evoke dendritic Ca^2+^ spikes by supra-threshold current injections to the trunk (Figure 3E).

**Figure 3.**
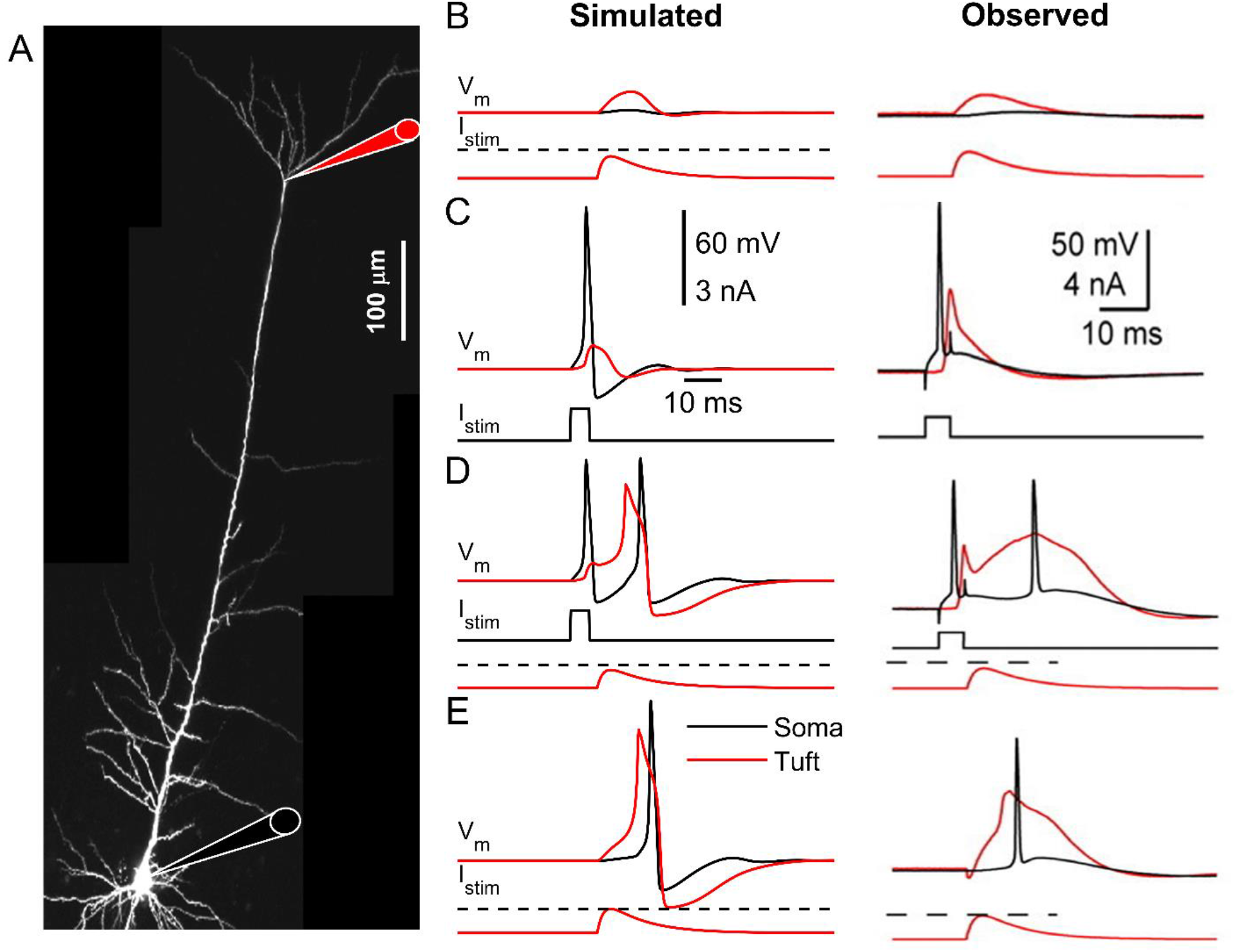
Back-propagating AP activated Ca^2+^-spike firing. **A** – Micrograph of a L5-PC with recording locations at the soma (*blue*) and distal trunk (*red*) indicated with a schematic pipette. **B** – Simulated (left) and observed (right, Schaefer et al., (2003)) subthreshold current injected into the apical dendrites creates only subthreshold somatic and dendritic depolarization. **C** – Simulated and observed suprathreshold somatic current pulse elicits an AP that propagates back to the apical dendrites creating a dendritic depolarization but no dendritic Ca^2+^-spike. **D** – Simulated and observed combined somatic and tuft stimulation evokes an AP, a dendritic Ca^2+^-spike, and another somatic AP following the onset of the dendritic Ca^2+^-spike. **E** – Simulated and observed suprathreshold stimulation of distal apical dendrites evokes a dendritic Ca^2+^-spike. Scales in C are common for all simulated (left) and observed (right) results. Red: apical-dendrite/trunks, Black: basal-dendrites/soma, and Dashed Line: dendritic threshold.

#### Critical frequency (CF) for Ca^2+^-spike generation

We next investigated the influence of the frequency of short somatic current stimulation on Ca^2+^-spikes occurrence. To that end, we simulated the soma stimulation with trains of brief supra-threshold pulses (2 *ms*) at different frequencies eliciting trains of somatic APs. As previously reported (Larkum et al., 1999a; Berger et al., 2003), only AP trains above a CF (149 *Hz* in the model) evoked Ca^2+^-spikes. Figure 4A illustrates the somatic and apical-dendrite/trunk responses to somatic stimulation below and at the CF. Figure 4B shows the intracellular Ca^2+^ concentration dynamics for both stimulation paradigms, which resemble experimental data (Larkum et al., 1999a).

**Figure 4.**
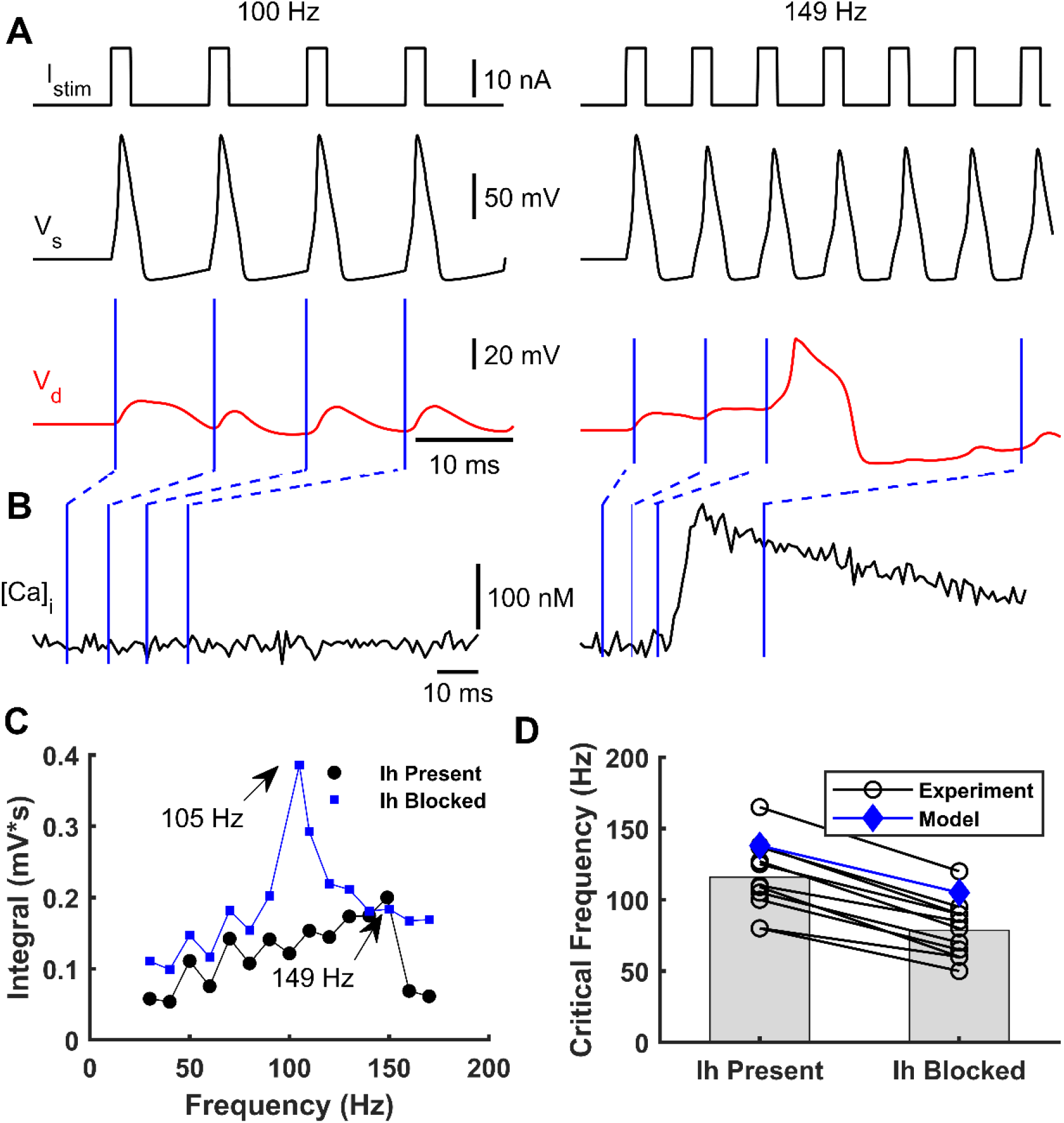
Effect of somatic stimulation frequency on dendritic Ca^2+^-spike occurrence. **A** – A simulated train of brief suprathreshold pulses at frequencies of 100 *Hz* (left) and 149 *Hz* (right) (top) was injected into soma eliciting a train of APs (*black*, below). Only the 149 *Hz* train evoked a dendritic Ca^2+^-spike (*red*, below). **B** – Intracellular dendritic Ca^2+^ concentration during somatic stimulation at 100 *Hz* (left) and 149 *Hz* (right). Blue lines indicate the Ca^2+^ concentration at each time instant of the dendritic voltage traces indicated in panel A. **C** – Integrated area below the dendritic voltage traces as a function of the AP frequency with (blue) and without (black) blocking the I_*h*_ current. CFs of 105 *Hz* and 149 *Hz* were obtained when I_*h*_ was present and absent, respectively. **D** – Observed shift in CF after blocking I_*h*_ for eleven cells (black circles, numerically estimated from Berger et al., (2003)) and simulated with the model (blue diamonds).

We also studied how the CF varied with the presence or absence of the I_*h*_ current in the distal apical dendrites. We simulated a L5-PC without I_*h*_ current at the apical-dendrite/trunk compartment responding to the same trains of supra-threshold currents at the soma with different frequencies. To quantify the CF, we measured the area below the dendritic voltage traces and plotted them as a function of AP frequency. When the I_*h*_ current was blocked relative to present, the CF was lower by about 40 *Hz* (Figure 4C). Furthermore, we compared the CF values with and without the I_*h*_ current predicted by our model with those predicted by experimental data from eleven L5-PCs (Berger et al., 2003) (Figure 4D). In both the observed and simulated data, the CF is reduced at least by 30-40 *Hz* when the I_*h*_ current is blocked. The CFs predicted by our model are slightly higher than the mean observed CFs, but they fell within the observed range.

#### Reproducing Ca^2+^-spike dependent local field potentials

To examine the capabilities of this minimal L5-PC model, we tested whether non-synaptic events such as Ca^2+^-spikes can be detected in the evoked LFPs as reported by Suzuki and Larkum, (2017). To that end, we simulated a collection of 1,000 model L5-PCs (Figure 1C). In the experimental paradigm, simultaneous stimulation of the soma of L5 PCs was achieved using an optogenetic approach (Suzuki and Larkum, 2017); hence, no synaptic connections were considered in our simulations. The simulated L5-PCs were stimulated with a noisy, 20 ms duration, current pulse with a mean amplitude that generates AP trains at a frequency above the CF. The mean input current was strong enough to generate somatic AP trains and therefore evoked dendritic Ca^2+^-spikes (Figure 5A). Figure 5B shows the raster plots and associated post-stimulus time histograms of 100 randomly selected L5-PCs (top), with the timing for typical Na^+^-APs and Ca^2+^-spikes. After the somatic stimulation ceased, somatic Na^+^-APs were only elicited because of the non-linear changes in the somatic-dendritic gain of these cells.

**Figure 5.**
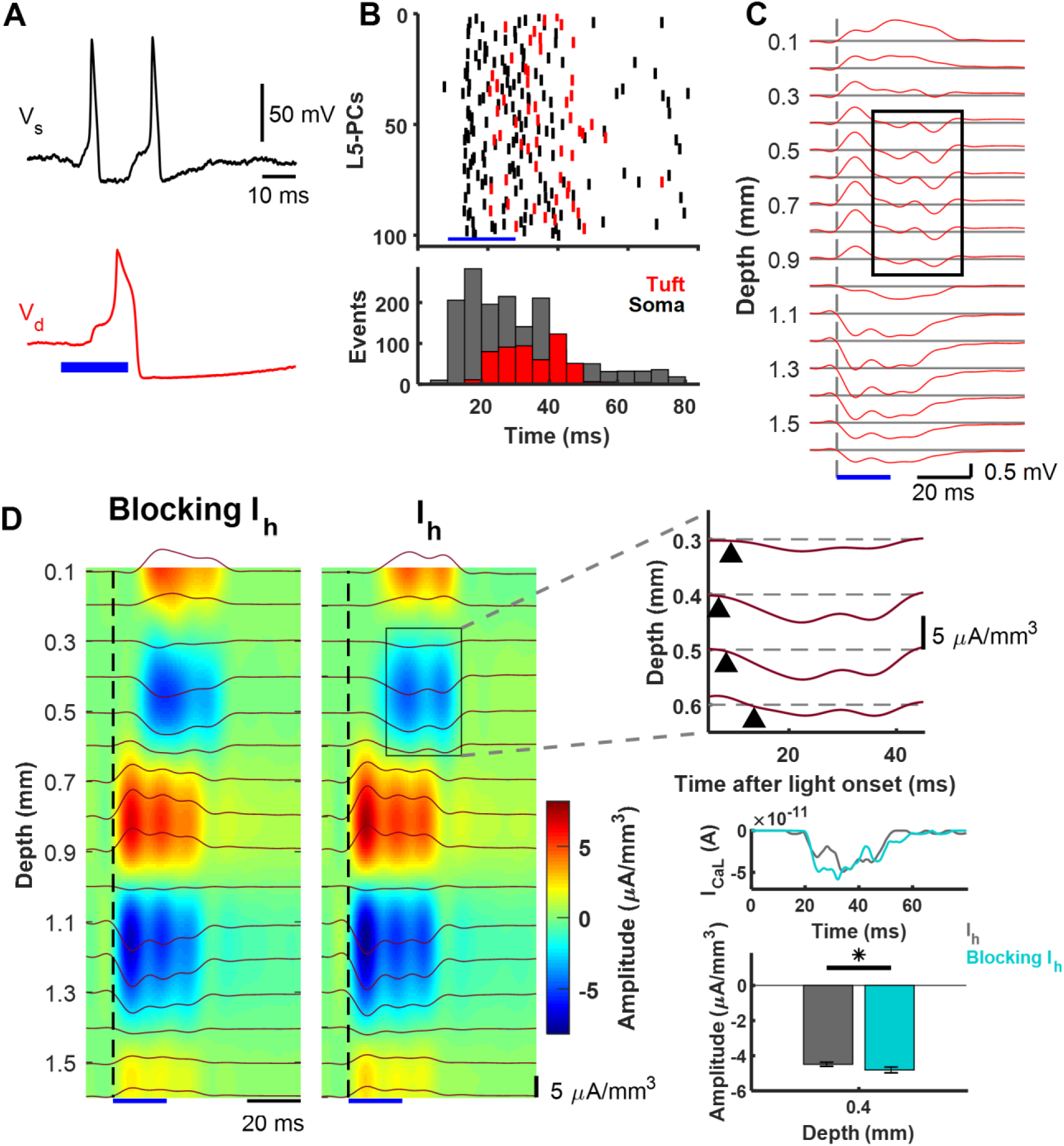
LFP and CSD derived from dendritic Ca^2+^-spikes in a collection of L5-PCs. **A** – Basal-dendritic/somatic (black, V_s_) and apical-dendritic/trunk (red, V_d_) simulated responses of a collection of 1,000 L5-PCs to suprathreshold optogenetic stimulation above the CF. **B** – Raster plots (top) and post-stimulus time histogram (bottom) of 100 randomly selected L5-PCs (top) showing spike times of Na^+^-APs (black) and Ca^2+^-spikes (red). The total number of Na^+^-AP and Ca^2+^-spike events every 5*ms* is shown (bar plots, bottom). **C** – LFPs evoked by the simulated optogenetic stimulation of the collection of L5 PCs calculated on an array of 16 microelectrodes (100 *μm* separation). Voltage traces at each depth are averaged over 10 simulated trials. The black rectangle indicates the delayed sink associated with the dendritic Ca^2+^-spike. **D** – CSD analysis of the evoked LFPs averaged over 10 trials without (left) and with (right) the I_*h*_ current. The top right panel expands the selected area to reveal the delayed sink associated with the Ca^2+^-spikes arising earliest 0.4 *mm* below the pia matter. Middle right panel plots averaged ***I*_*CaL*_** current in the trunk of the L5-PCs, showing an amplitude increase 25-35 *ms* after stimulation when I_*h*_ was blocked. Lower right panel compares the average with SEM of the amplitude of this current sink with and without blocking the I_*h*_ current, which was significantly different (p = 0.0089, Wilcoxon signed-rank test, N = 10 trials). The blue bar in all the plots indicates the time window for the optogenetic stimulation.

Figure 5C illustrates the averaged LFPs evoked by optogenetic stimulation of the collection over 10 trials. We observed an early sink between 1.0-1.3 *mm* below the pia matter, which was accompanied by two sources, one stronger between 0.7-0.9 *mm* and another weaker between 1.4-1.6 *mm.* According to our model, the sink was caused by large *I*_*Na*_ inward currents at the level of the axon hillock due to the optogenetically induced APs. The two sources were caused by a combination of *I*_*Kdr*_ and the returning capacitive/leak outward currents through the top soma-oblique dendrites and the basal dendrites. The relative amplitudes of these two sources can be adjusted by means of parameters *α*_*Kdr*_ and *α*_*C*_. To create **Figure 5**, these parameters were set to 0.5 and 1/3, respectively. We also observed a 20-30 *ms* delayed sink between 0.3-0.6 *mm* below the pia matter, which was accompanied by a very superficial (0.1-0.2 *mm*) source, also delayed. This late sink appeared during the same interval in which the collection of L5 PCs generated more Ca^2+^-spikes (Figure 5B bottom). Hence, we believe it was caused by the *I*_*CaL*_ inward current.

According to our model, the superficial sources resulted from a combination of *I*_*M*_, *I*_*Ks*_, and the returning capacitive/leak outward currents through the apical-dendrites. Because of its reversal potential, the cation current *I*_*h*_ could be either a source or a sink during a Ca^2+^-spike at a very superficial level. The relative amplitude of this delayed sink-source was adjusted using parameter *α*_*Ks*_ = 1 to reproduce a similar CSD pattern as that reported by Suzuki and Larkum (2017). The CSD analysis clearly revealed the presence of such a sink/source current density distribution (Figure 5D, right color map panel (I_*h*_) and expanded plot, respectively). Since we did not consider synaptic connections between the L5-PCs, the above results suggest that the late sink is associated with the dendritic Ca^2+^-spikes.

Finally, we investigated the influence of the I_*h*_ current on the source-sink pattern generated by the dendritic Ca^2+^-spikes. We repeated the simulations, but now without the I_*h*_ current in the apical-dendritic compartment (Figure 5D). In agreement with the experimental data (Suzuki and Larkum, 2017), we found that the amplitude of the delayed sink in layer 2/3 is significantly increased by blocking the I_*h*_ current (p = 0.0089, Wilcoxon signed-rank test, N = 10 trials). Since the superficial source in Suzuki and Larkum (2017) was very close to the pia boundary, we believe the iCSD method used by the authors might have misestimated this source. Therefore, we did not compare the experimental effect of blocking I_*h*_ on that superficial sources with that predicted by our model. The cation current I_*h*_ was too small in amplitude to produce any detectable change in the CSD when blocked. However, this current was crucial to produce a shifted resting membrane potential of +10 mV (Figure 1C) in the apical-dendrite/trunk compartment, which disappeared when I_*h*_ was blocked. As the trunk resting membrane potential became more negative, the effect of *I*_*CaL*_ was larger between 25-35 *ms* after stimulation (**Figure 5**D), producing a more intense delayed sink during Ca^2+^-spiking at the level of the L5-PC trunk.

## Discussion

Synaptic integration in apical dendrites of L5-PCs is facilitated by several unique characteristics of these neurons: **(i)** the f-I curves with differentiated sensitivity for the soma and distal trunk, **(ii)** the BAC firing-based amplification of coincident apical-dendritic inputs, **(iii)** the CF effect for eliciting Ca^2+^-spikes in the distal trunk, and **(iv)** its related increases in intracellular [*Ca*^2+^]_*i*_ in apical-dendrites strengthening NMDA synaptic efficacy. Biophysical models with different level of complexity have been proposed to account for single L5-PC features (Rapp et al., 1996; Larkum et al., 2004; Chua et al., 2015; Yi et al., 2017) or combinations of features (Schaefer et al., 2003; Hay et al., 2011; Bahl et al., 2012; Almog and Korngreen, 2014; Mäki-Marttunen et al., 2018). Models explaining combinations of features require many compartments to capture realistic L5-PC morphology and more than four ionic channels per compartment, which substantially increase the computational cost and time (Table 3). Consequently, using these models in large-scale simulations of collections of L5-PCs requires special computational resources and extensive time. Moreover, fitting them to large-scale electrophysiological data (e.g., LFP and EEG) will be challenging, limiting interpretability and applications to other research areas. Also, no previous model has accounted for the shift in the CF due to the influence of I_*h*_, explained the CSD patterns associated with dendritic Ca^2+^-spikes evoked by somatic stimulation of PCs above the CF, or the effect on these patterns of blocking I_*h*_ (Suzuki and Larkum, 2017). We proposed a 2-compartment model of L5-PCs with just seven ion channels that explain qualitatively all abovementioned features.

### Layer 2/3 PCs vs. layer 5 PCs

Does our model account for characteristics of PCs in layer 2/3 (L2/3)? Even though they share many electrophysiological properties, L2/3-PCs have distinct features that differentiate their role in the cortical microcircuit (Larkum et al., 2007). Similar to L5-PCs, L2/3-PCs have excitability zones in both the axon initial segment and the distal apical dendrites. These act as two different functional compartments that allow these neurons to associate inputs coming to the distal apical dendrites with those coming to the soma or basal dendrites. Large Ca^2+^ influx and regenerative dendritic potentials are also evoked by back-propagating action potentials above a CF. Moreover, as in L5-PCs (Pérez-Garci et al., 2006), GABAergic inhibitory inputs to the distal apical dendrites cause long-lasting reduction of dendritic activity. However, unlike L5-PCs, L2/3-PCs do not show long plateau-like dendritic depolarizations. Consequently, brief dendritic spikes have less influence on somatic AP output. In fact, isolated dendritic potentials in response to supra-threshold dendritic stimulation are more common than dendritic spikes coupled to a somatic AP. Furthermore, though coincident inputs to both functional compartments reduce the threshold for dendritic spike generation, stronger dendritic inputs are needed to evoke an extra somatic AP. In addition, L2/3-PCs display little attenuation in the dendritic response to long current injections suggesting a low density of I_*h*_ channels in the dendrites, described as sag by Larkum et al. (2007).

### Functional Implications: Microcircuitry underlying cognitive control

Cognitive control involves the suppression of automatic or impulsive behavior for successful goal-directed behavior. Some models of cognitive control formalize this function as the co-activation of two conflicting action plans, which need to be resolved for correct performance (Botvinick et al., 2001). Coincidence detection can also support *error detection* – a mismatch (or conflict) between task goals and actual behavior – and *prediction error* – a mismatch between expected and experienced outcomes (Alexander and Brown, 2011; Bastos et al., 2012; Cohen, 2014). Human and macaque electrophysiology experiments have characterized a negativity in scalp potentials associated with these cognitive functions (Gehring et al., 1993). Two components: an N2 for conflict detection and the ERN for error detection. While the N2 and ERN are indices of cognitive control, studying signal processing at the microcircuit level is essential to understanding actual mechanisms (Cohen, 2017). Our biophysical model offers a powerful tool to test different hypotheses and instantiate circuit models motivated by recent research sampling neural spiking and LFP across frontal cortical layers (Chandrasekaran et al., 2017; Bastos et al., 2018; Sajad et al., 2019).

Recent models have proposed that conflict detection can be achieved by the detection of coincident synaptic inputs in the medial frontal cortex (Alexander and Brown, 2011; Cohen, 2014; Dembrow et al., 2015). L5-PCs can provide the neural substrate for the coincidence detection as they have large dendritic trees that allow for integration across inputs from cognitive, limbic, and motor structures (Huerta and Kaas, 1990; London and Häusser, 2005; Morecraft et al., 2012; Beul and Hilgetag, 2015). Recently, we found that following errors in the stop-signal task, in a medial frontal area, error-related neural spiking was first observed in putative pyramidal neurons in L5 and lower L3 (Sajad et al., 2019) concurrent with sinks in current in the superficial layers where these neurons extend their dendrites (Sajad et al., 2017). Figure 6 diagrams our conjecture on the role of L5-PCs in error detection in agranular cortex guided by the knowledge of the microcircuitry and known anatomical connections. L5-PCs can detect the coincidence of an efferent copy of the motor command from the mediodorsal thalamus and the task rule from prefrontal cortex (Sajad et al., 2019). A mismatch between the two signals can result in spiking activity that can project extrinsically to other structures (Barbas, 2015) and intrinsically to other neurons in the microcircuit (Douglas et al., 1995; Haeusler and Maass, 2007; Kajikawa and Schroeder, 2011). L5-PCs are densely interconnected with each other (not shown) resulting in rapid synchronous excitation of a large number of L5-PCs upon receiving input currents (Hempel et al., 2000; Wang et al., 2006; Morecraft et al., 2012). L5-PCs are also connected to inhibitory interneurons, which result in the control of this excitation. Noteworthy, recent evidence suggests inhibitory neurons in agranular cortex support more intra-than inter-laminar connections (Kätzel et al., 2011; Beul and Hilgetag, 2015). Hence, the inter-laminar inhibitory projections depicted in Figure 6 represent inhibitory influences that are likely mediated by additional PCs and interneurons in L3 and L5 (not shown).

Previous studies have suggested that the patterns of interconnectivity of PCs and their inhibition, particularly by the somatostatin-positive neurons, is important for controlled patterns of theta-band rhythmogenesis in the medial frontal cortex (Mainen and Sejnowski, 1996; Cohen, 2014), and its genesis has been associated with the role of the I_*h*_ current (Ulrich, 2002). Also, intrinsic signal processing, involving inhibition of L5-PCs by interneurons in the upper layers has been associated with the generation of gamma oscillations (Buzsáki and Wang, 2012; Bastos et al., 2018). These oscillations result in scalp potential reflections of the gamma rhythm as well as the theta rhythm, one the hallmarks of error and conflict detection (Tallon-Baudry and Bertrand, 1999; Cohen and Donner, 2013).

**Figure 6.**
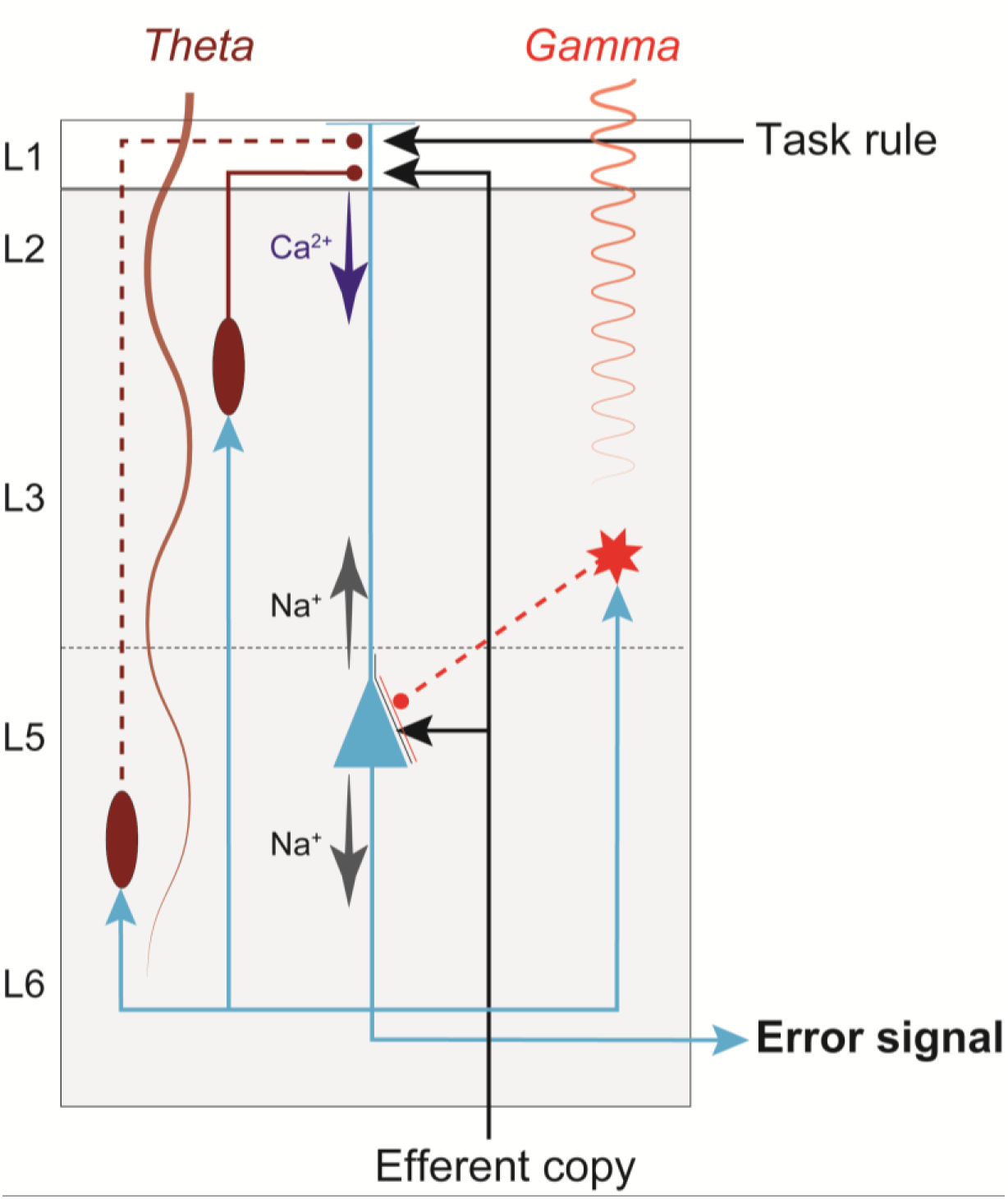
Cortical microcircuit for coincidence detection underlying cognitive control. The simplified diagram of circuitry embedding a L5-PC (blue) in agranular cortex with soma (triangle) located in L5, dendrites that extend up to L1, and axons (blue arrows) that project both intrinsically, innervating inhibitory neurons (red) and other pyramidal neurons (not shown) in the microcircuit, and extrinsically, innervating other brain areas. This figure illustrates how dendritic dynamics can contribute to an error signal. An efferent copy of a motor command is delivered through a feedforward thalamic pathway, terminating on the L5-PC soma and apical dendrite. A task rule signal from prefrontal cortex is delivered through a feedback pathway, terminating on the L5-PC apical dendrites. The soma of a L5-PC (blue triangle) generates Na^+^-APs that propagate intracortically to Martinotti cells (ovals) and other inhibitory interneurons (star). The Martinotti cells terminate on the L5-PC apical dendrites, while the other interneuron terminates on the soma. Note that because inhibitory neurons in agranular cortex largely make intra-laminar projections, the inter-laminar inhibitory projections depicted here (dashed red lines) represent connections that are likely mediated by additional PCs and interneurons in L3 and L5 (not shown). The dynamics of this connectivity induces Ca2+ spikes, which amplify the coincidence of the efferent copy and the task rule to generate an error signal. These neuronal events are signaled by the generation of theta band LFP from deeper layers and gamma band LFP from superficial layers (indicated by labeled oscillations).

While Figure 6 provides one explanation for signal flow within the microcircuit, it is far from complete and relies on untested assumptions. For instance, the location where inputs to L5-PCs converge and the mechanism for how these signals are integrated at the biophysical level remains technically challenging to study (Stuart and Spruston, 2015). Furthermore, the interaction between L5-PCs and other neurons in the microcircuit remains unclear. The biophysical model proposed in the current study provides an essential tool for further testing between and refining competing hypotheses. Future work needs also the development of similar biophysical models for L2/3-PCs and other neurons in the microcircuitry.

### A necessary step towards understanding the origin of the ERN

The proposed model will be useful for another research goal of developing a forward model of the ERN component. Clearly, the EEG arises from the activity of neurons in the brain tissue, but the detailed relationship to activity within neocortex remains unclear (Riera et al., 2012; Einevoll et al., 2013; Reimann et al., 2013). Recently, we have shown that error-related spiking activity of neurons in the upper layers, but not lower layers of monkey supplementary eye field predicts the magnitude of the ERN (Sajad et al., 2019). Also, recent work recording from single neurons in humans have shown coupling between error neuron activity and intracranial EEG (Fu et al., 2019). Somatic action potentials are unlikely to directly influence the EEG due to their voltage dynamics; however, bursts of action potentials from a large population of neurons can influence the EEG (Buzsáki et al., 2012). Furthermore, the large-scale EEG topography of Ca^2+^-spikes remains unknown (Suzuki and Larkum, 2017). The proposed biophysical model of L5-PC can be used to directly examine how current flow resulting from neuronal activity within the microcircuit, down to fine details of current dynamics, can result in LFP and scalp EEG (Riera et al., 2012). Establishing a link between specific microcircuit motifs and fluctuations in the scalp potentials can render ERN more effective markers of specific cortical processes and stronger diagnostic tools for patients with compromised cognitive control functions.

## Acknowledgments

This work was supported by FIU SEED – grant Wallace Coulter Foundation (BH, JR), R01-EY019882 (JDS, GFW, BH, JR), and by Robin and Richard Patton through the E. Bronson Ingram Chair in Neuroscience (JDS). The authors thank Michelle Schall and Arash Moshkforoush for helpful discussions and comments on the manuscript.

